# SARS -CoV-2 T-cell immunity to variants of concern following vaccination

**DOI:** 10.1101/2021.05.03.442455

**Authors:** Kathleen M.E. Gallagher, Mark B. Leick, Rebecca C. Larson, Trisha R. Berger, Katelin Katsis, Jennifer Y. Yam, Gabrielle Brini, Korneel Grauwet, MGH COVID-19 Collection & Processing Team, Marcela V. Maus

## Abstract

Recently, two mRNA vaccines to severe acute respiratory syndrome coronavirus 2 (SARS-CoV-2) have become available, but there is also an emergence of SARS-CoV-2 variants with increased transmissibility and virulence^1–6^. A major concern is whether the available vaccines will be equally effective against these variants. The vaccines are designed to induce an immune response against the SARS-CoV-2 spike protein^7, 8^, which is required for viral entry to host cells^9^. Immunity to SARS-CoV-2 is often evaluated by antibody production, while less is known about the T-cell response. Here we developed, characterized, and implemented two standardized, functional assays to measure T-cell immunity to SARS-CoV-2 in uninfected, convalescent, and vaccinated individuals. We found that vaccinated individuals had robust T-cell responses to the wild type spike and nucleocapsid proteins, even more so than convalescent patients. We also found detectable but diminished T-cell responses to spike variants (B.1.1.7, B.1.351, and B.1.1.248) among vaccinated but otherwise healthy donors. Since decreases in antibody neutralization have also been observed with some variants^10–12^, investigation into the T-cell response to these variants as an alternative means of viral control is imperative. Standardized measurements of T-cell responses to SARS-CoV-2 are feasible and can be easily adjusted to determine changes in response to variants.

## INTRODUCTION

Most reports of immunity to SARS-CoV-2 infection or vaccination focus on humoral immunity by quantifying anti-spike IgG, IgM, or IgA levels in peripheral blood^13–17^. Antibodies to the SARS-CoV-2 spike protein can be detected in convalescent individuals following natural infection and correlate with virus neutralizing activity^13, 14, 18^. In individuals who develop antibodies, some reports show that IgG levels wane over the first 3–6 months^13, 19–22^, despite the persistence of memory B cells^13^, whereas others have demonstrated a more sustained response 5–8 months following infection^18, 23, 24^. Continued immunity in individuals is imperative for developing herd immunity and preventing virus spread.

In addition to humoral immunity, T-cell immunity is important for eliminating infected cells and promoting antibody class switching. The development of memory T-cells recognizing SARS-CoV-2 is likely to be important for long-term protection. With other closely-related coronavirus infections, antibody titers were initially weak or diminished within 2–3 years^25–28^. In contrast, virus-specific T-cells, which can develop in the absence of seroconversion^29^, have been detected for 6–11 years^30, 31^. Similarly, SARS-CoV-2 specific T-cells are present in COVID-19 convalescent individuals^32–34^, even in the absence of seroconversion^32^ or symptomatic infection^34^. This suggests that cellular immunity may play a larger role in sustained protection and a subset of the population that does not have detectable antibodies may nonetheless be protected due to T-cell mediated immunity.

## RESULTS

### SARS-CoV-2 ELISpot T-cell immunity test

To measure SARS-CoV-2 T-cell immunity in convalescent and vaccinated individuals, we developed an IFNγ ELISpot assay that quantifies IFNγ-producing T-cells in response to viral peptides. We used commercially available peptide pools from the initial SARS-CoV-2 spike and nucleocapsid protein sequences identified in Wuhan^35^. Given its size, the spike protein was split between two pools, A and B, with A containing amino acids (AA) 1–643 and B containing AA633– 1273 (**Figure 1A**). We stimulated freshly isolated peripheral blood mononuclear cells (PBMC) with the peptide pools and quantified the median spot forming units (SFU) per 2.5x10^5^ PBMC from duplicate wells (**Figure 1B**).

**Figure 1.**
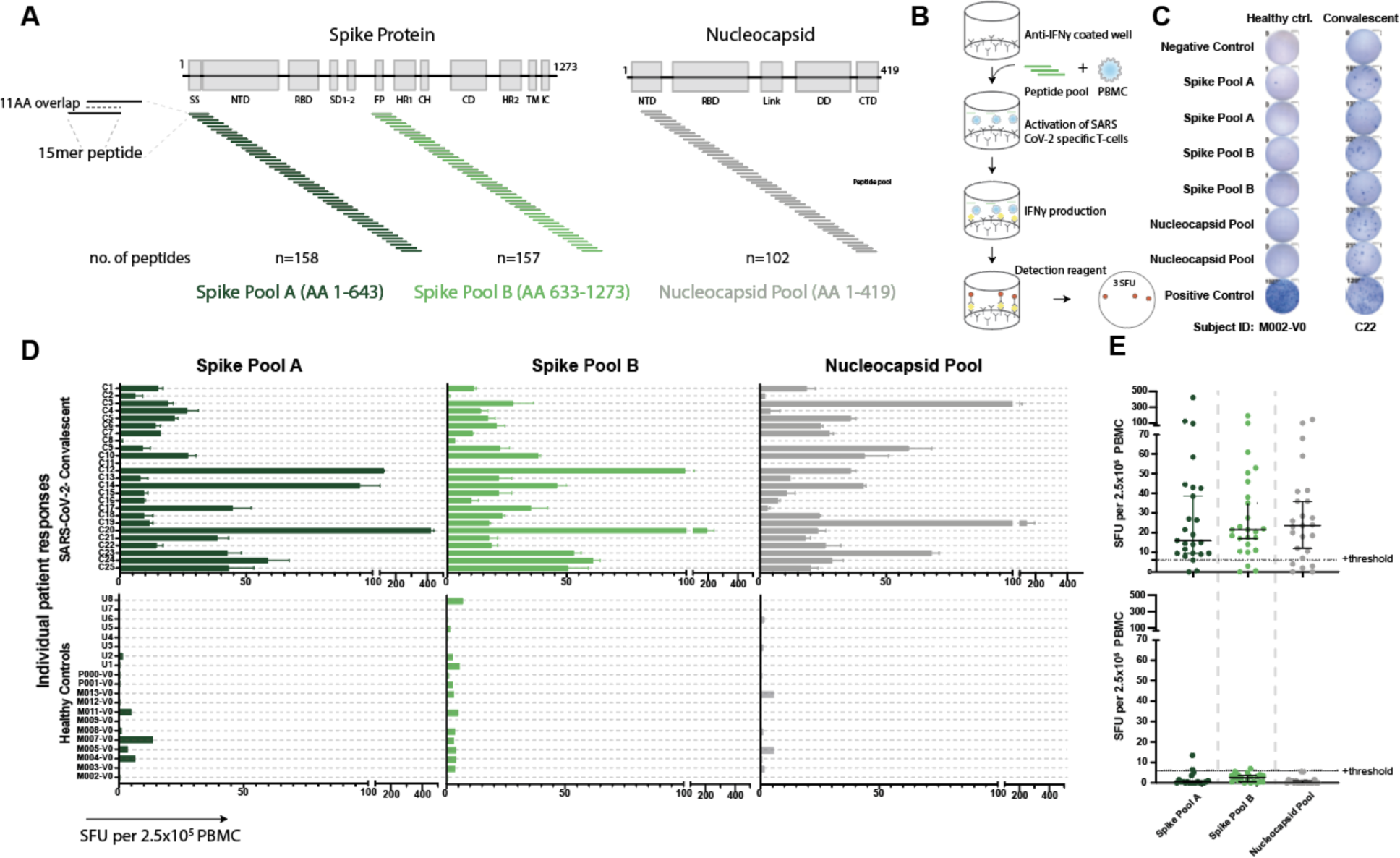
Validated IFNγ ELISpot assay discriminates between SARS-CoV-2 naïve and convalescent T-cell responses. **A.** SARS-CoV-2 spike and nucleocapsid protein domains with corresponding peptide pools. **B.** Schematic of IFNγ ELISpot assays. **C**. Representative ELISpot wells from a healthy control (M002) and a SARS-CoV-2-convalescent donor (C22) with the number of spot forming units (SFU) per 2.5x10^5^ PBMC quantified (number next to well). **D**. SFU are shown for each subject in the SARS-CoV-2 convalescent (n=25) or healthy control (n=19) groups. Bars represent mean ± standard error of the mean. **E**. Composite ELISpot results from D. Lines represent median ± 95% confidence intervals. Dotted line indicates the positive threshold of 6 SFU per 2.5x10^5^ PBMC.

To establish the SFU threshold for the peptide pools and the sensitivity and specificity of the assay, we first evaluated responses in two donor groups: healthy controls who had no known history of SARS-CoV-2 infection or definitive exposure as defined by the CDC (n=19; **Extended Data Table 1**; 13 of the 19 had serum available for antibody testing and were negative for anti-spike-IgG/A/M), and a group of convalescent patients who had recovered from confirmed COVID- 19 (n=25; **Extended Data Table 2;** 2 of the 25 were positive by antibody testing to spike protein but had tested negative by PCR). The median time from diagnosis to ELISpot assay was 162 days (range of 83–237 days; **Extended Data Figure 1**). The healthy control cohort had low background responses to all SARS-CoV-2 peptide pools, compared to strong T-cell responses in the convalescent cohort (**Figure 1C, D**). Convalescent patients had a median SFU of 16.0 for spike pool A, 21.5 for spike pool B and 23.5 for the nucleocapsid pool, compared to ≤2.5 SFU in the healthy controls (**Figure 1E**). There was no correlation between SFU and the days from diagnosis or patient age (**Extended Data Figure 2**). Only one convalescent donor (C11) had no response to any of the peptide pools. Interestingly, this individual had tested positive for SARS-CoV-2 by PCR but tested negative for antibodies to spike protein despite having mild symtpoms.

**Figure 2.**
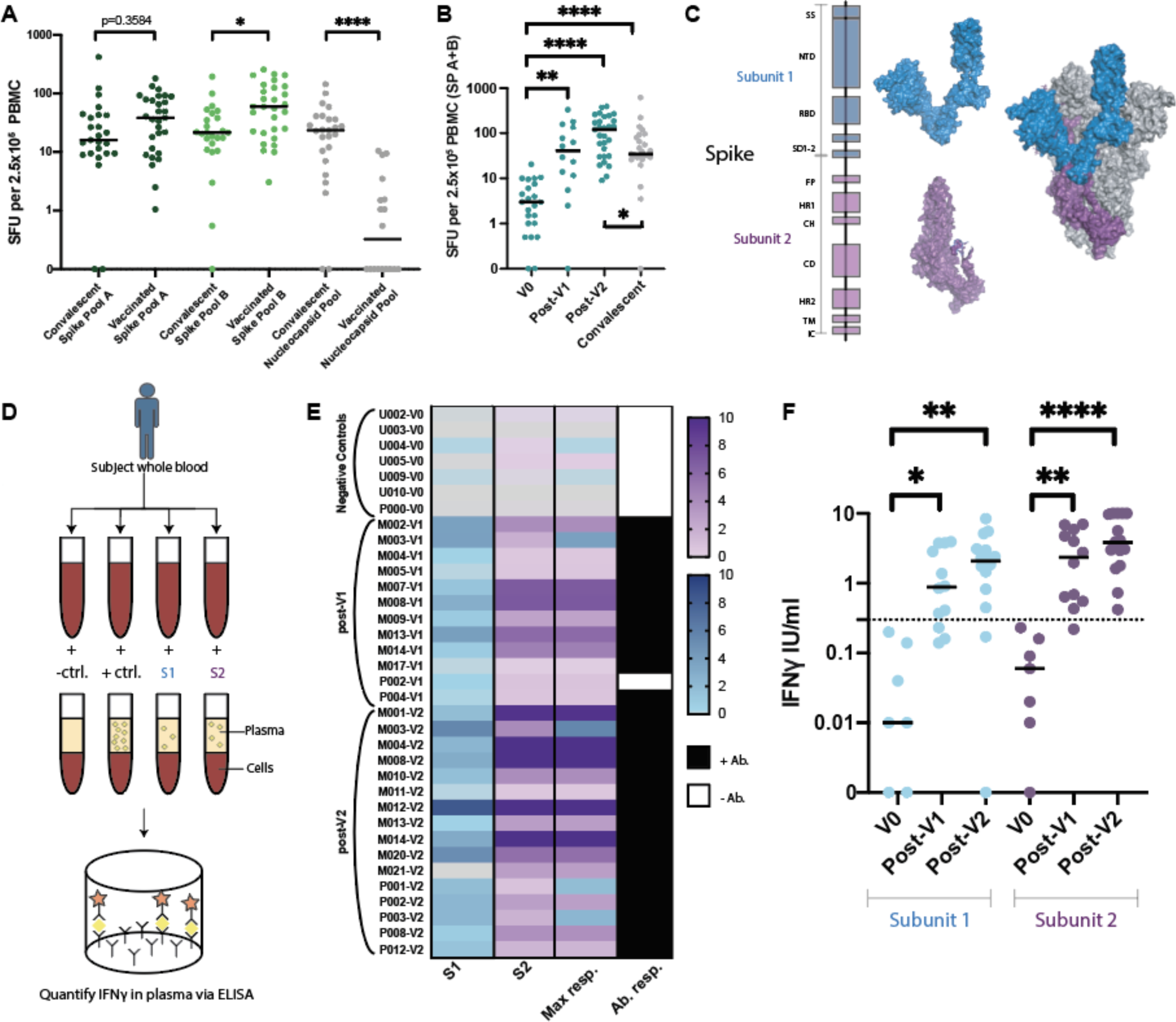
SARS-CoV-2 specific T-cell responses in vaccinated cohorts. **A.** ELISpot results for convalescent patients compared to fully vaccinated donors (V2) (Kruskall Wallis with Dunn’s multiple comparison test where *p=0.0311 and *****p<0.00001). **B.** ELISpot results assessed prior to vaccination (V0), post first vaccination dose (V1), and post second vaccination dose (V2) (Kruskal-Wallis with Dunn’s multiple comparisons test where, ****p<0.0001, **p=0.018). Differences between convalescent results versus V0 where ****p<0.0001, *p=0.0219 (Mann-Whitney test). **C.** Spike subunit 1 (S1) and subunit 2 (S2) structures shown independently (left) and as a representative monomer of the *in situ* trimer (right, two additional monomers shown in gray). **D.** Schematic of novel whole blood IFNγ release assay (hereon referred to as “whole blood assay”). **E.** Heatmap of IFNγ release (IU/ml) to S1 and S2 subdomains in the whole blood assay from donors for which all results were available. The highest response to either S1 or S2 is indicated in Max response column. Paired serums for anti-spike IgG/IgA/IgM reported as positive (black) or negative (white). **F.** Composite whole blood assay IFNγ responses (Kruskal-Wallis test with Dunn’s multiple comparisons test where S1: *p=0.0461, **p=0.0028 and for S2: **p=0.034 and ****p<0.0001). Dotted line indicates a positive threshold of 0.3 IU/ml. Median shown for A, B, and F. SP= Spike pool.

Using these results, we performed receiver operator characteristic (ROC) curve analyses for the individual peptide pools (**Extended Data Figure 3A**). The relationship between the average SFU in response to either spike pool was evaluated for the associated sensitivity and specificity (**Extended Data Figure 3B**). A positive threshold of 6 SFU provided the optimal balance between sensitivity (92.0%) and specificity (90.0%). Using the same positive threshold of 6 SFU, the nucleocapsid pool response had a sensitivity of 80% and a specificity of 95%. This assay represents a unified and standardized approach to measure T-cell responses to SARS-CoV-2 that could be used by clinical labs with experience in performing ELISpot tests.

**Figure 3.**
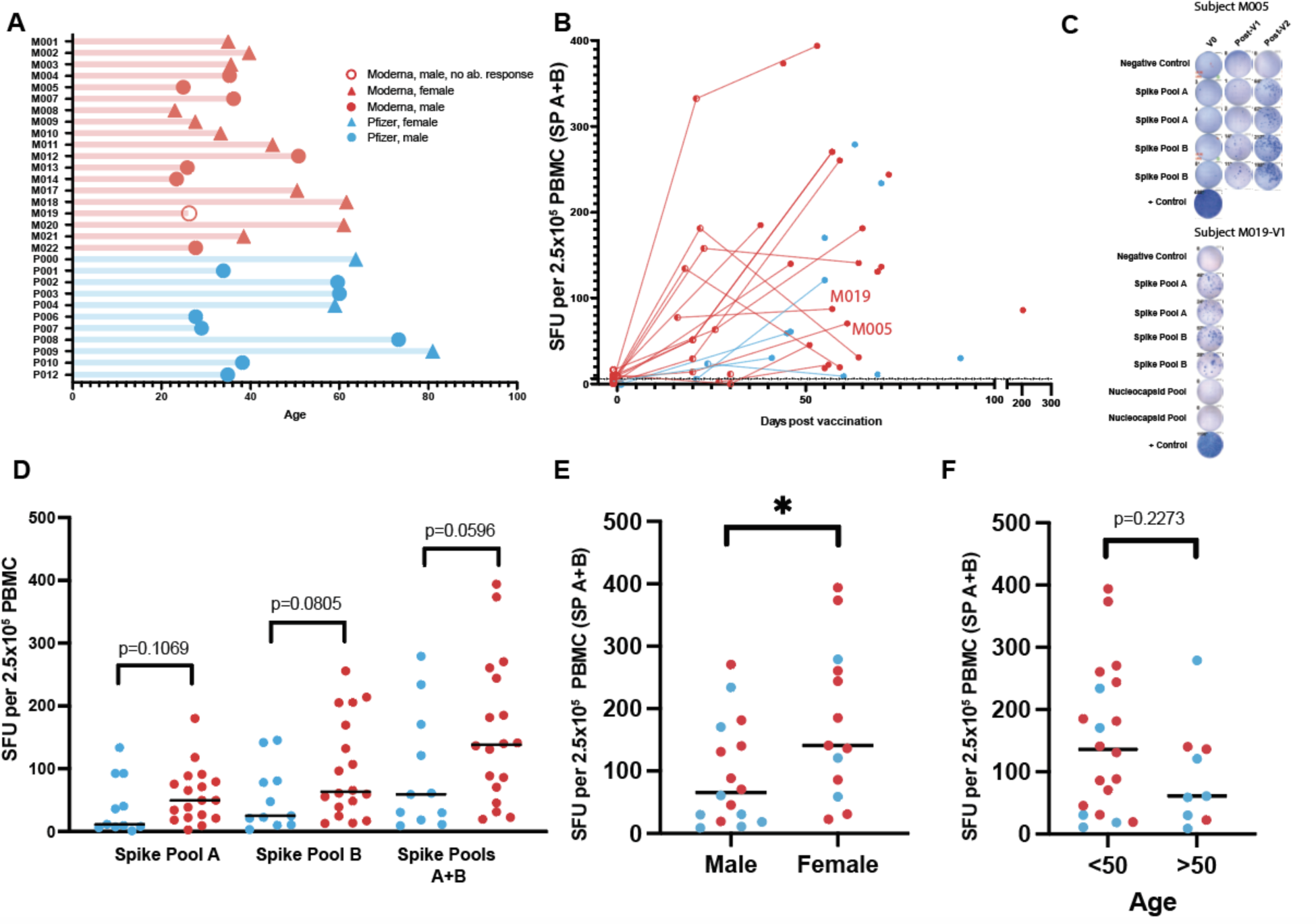
Magnitude of the T-cell response in vaccinated individuals. **A.** Age of vaccinated donors assayed for ELISpot responses (median=35.9 years). Vaccine: Moderna (n=19), Pfizer (n=11). Sex: male (n=16), female (n=14). Seroconversion: Positive antibody response (n=29), Negative (n=1). **B.** Responses from spike pools (A+B) versus assay date in relation to first vaccine dose 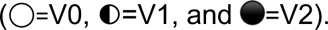. Days from V1 median=22; range=16-31. Days from V2 median=59, range=38-204. **C.** Representative ELISpots for 2 donors shown. **D.** V2 ELISpot responses stratified by vaccine (unpaired t-test). **E.** V2 ELISpot responses stratified by sex (unpaired t-test *p=0.0411). **F.** V2 ELISpot responses stratified by age (unpaired t-test p=0.2273). Median shown for D-F.

### T-cell immunity following vaccination

As of January 2021, there were two vaccines with FDA emergency use approval in the United States. Both vaccines (developed by Pfizer/BioNTech and Moderna) deliver mRNA coding for the full-length spike protein, require two doses a minimum of 21 or 28 days apart, and provide 95% and 94.1% efficacy at preventing symptomatic COVID-19 infection respectively^7, 8^. We used our ELISpot assay to evaluate SARS-CoV-2 T-cell responses in vaccinated individuals prior to inoculation (V0/negative controls; median 10.5, range 0–156 days prior to injection), after the first dose (post-V1; median 22 days, range 16–30 days), and after the second dose (post-V2; median 59 days after the initial dose, range 38–204 days). PBMC from 29 donors post-V2 (11 Pfizer, 18 Moderna) were tested using the IFNγ ELISpot assay. This cohort of vaccinated donors had no history of COVID-19 symptoms or positive SARS-CoV-2 test prior to vaccination.

Similar to results from the phase I/II vaccine trials^36, 37^, all fully vaccinated donors developed positive T-cell responses to one or both spike peptide pools. The median responses were higher in the fully vaccinated cohort compared to the convalescent cohort (**Figure 2A**). While, on average, convalescent patients were tested further from diagnosis than vaccinated donors from injection, no association was seen between days from positive test and ELISpot response (**Extended Data Figure 2A**). Interestingly, the magnitude of T cell response to spike pool B was higher than to pool A in vaccinated donors (p=0.0311), but not convalescent patients. T-cell responses against the nucleocapsid protein, which is not encoded in the SARS-CoV-2 vaccines, were lower in vaccinated subjects compared to the convalescent cohort (p<0.0001). Thus, positive responses (≥6 SFU/2.5x10^6^ PBMC) to the nucleocapsid protein likely reflect natural infection.

A portion of donors (n=14) were also tested post-V1, and T-cell responses were positive against at least one spike pool in 11/14 tested. The median SFU in response to spike pools A+B increased post-V1 (p=0.018) and post-V2, (p<0.0001) compared to baseline (**Figure 2B**). The cumulative SFU response to spike pool A+B post-V1 were of similar magnitude to those observed in the convalescent cohort, but were increased compared to the convalescent cohort post-V2 (p=0.0219; **Figure 2B**)

### High throughput whole blood T-cell assay

We next sought to develop a more straightforward assay for evaluating T-cell responses to vaccination that would be better suited for high-throughput processing, which would avoid the personnel and blood-processing requirements of ELISpot analysis and expensive peptide pools. To achieve this, we designed a whole-blood assay based on the *in vitro* diagnostic QuantiFERON-TB Gold Plus assay that is used to evaluate T-cell recognition of mycobacterium tuberculosis (TB) antigens when assessing latent TB infection. In our assay we focused on assessing T-cell responses to the spike protein, which consists of two main subdomains, S1 and S2, that can form a trimeric tertiary structure (**Figure 2C**). S1 contains the receptor-binding domain (RBD), which interacts with angiotensin-converting enzyme 2 (ACE2) on host cells^38, 39^, while S2 promotes fusion with the host cell membrane. We stimulated heparinized whole blood with positive and negative controls and with the individual monomeric forms of SARS-CoV-2 S1 or S2 proteins overnight and then measured IFNγ (IU/mL) released into the plasma using a QuantiFERON ELISA (**Figure 2D**). We used results from 7 unvaccinated/unexposed healthy controls and 16 vaccinated donors post-V2 (5 Pfizer, 11 Moderna; median 56 days following the first dose, range 38–204 days) to determine the positive threshold of the assay. Individual ROC curves for S1 and S2 were calculated (**Extended Data Figure 4A**). We then evaluated how assay sensitivity/specificity were impacted by altering the positive threshold when the response to both spike subdomains were combined (i.e. if the response to either S1 or S2 was greater than a given threshold value). A positive threshold of ≥0.3 IU/mL was selected to optimally discriminate between the two groups, with a sensitivity and specificity of 100% (**Extended Data Figure 4B**). This threshold value is similar to the positive cut-off of 0.35 IU/mL that is used in the *in vitro* diagnostic QuantiFERON-TB Gold Plus assay^40^.

**Figure 4.**
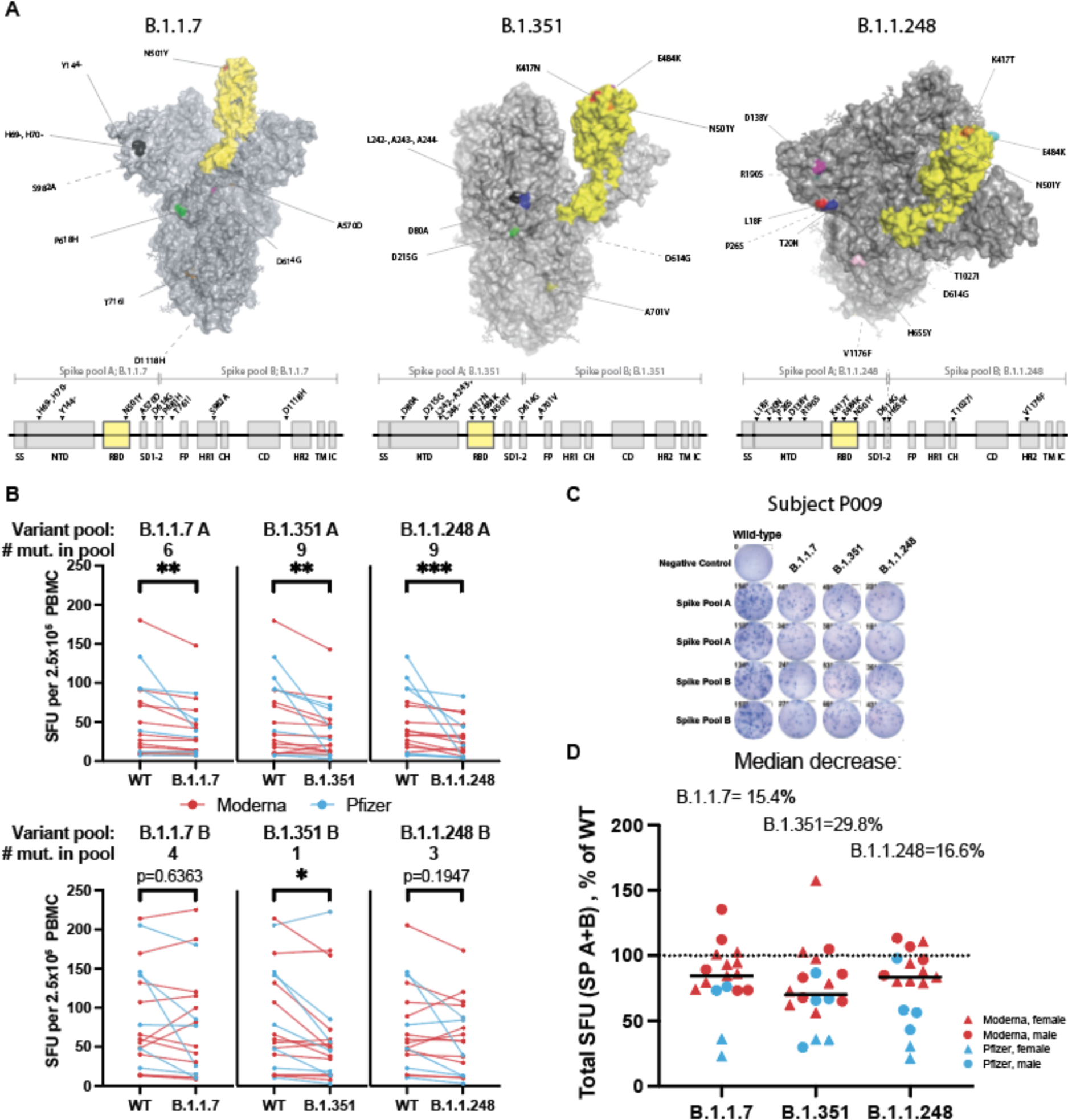
Spike variants B.1.1.7, B.1.351, and B.1.1.248 induce a diminished T-cell response compared to the wild type protein. **A.** SARS-CoV-2 spike protein variants B.1.1.7, B.1.351 and B.1.1.248 with mutations compared to the vaccine spike sequence noted (RBD in shown yellow). Location of mutations in primary sequence are depicted with reference to the corresponding spike pool affected. **B.** ELISpot response to mutant versus wild type (WT) spike peptide pools (n=18-20; Moderna=red, Pfizer=blue; Wilcoxon matched pairs signed-ranks test with Bonferroni correction; B.1.1.7 A **p=0.0015, B.1.351 A **p=0.0015, B.1.1.248 A ***p=0.0006; B.1.351 B *p=0.0144. **C.** ELISpot results against WT and variant peptides shown for a representative patient. **D.** ELISpot response to variant peptide pools relative to individual subject’s WT response. Medians shown as black lines. Dotted line represents response to WT (100%). Only subjects with a positive ELISpot against WT (≥6) were included for variant peptide analysis.

When visualizing IFNγ produced in response to S1 or S2, there was a clear distinction between the unvaccinated/unexposed healthy controls (V0) and vaccinated donors post-V2 (**Figure 2E**). All post-V2 donor samples had positive responses to either S1 and/or S2, with S1 responses lower compared to S2 (p=0.0063). Some donors (n=12) were also tested post-V1 (2 Pfizer, 10 Moderna; median 22 days post vaccination, range 18–30; **Figure 2E**). Only one donor (M017) had a negative result. This individual was also negative in the ELISpot assay yet had detectable anti-spike antibodies by IgG/IgA/IgM ELISA, and subsequently developed a spike response in the ELISpot assay post-V2, but could not be reassessed in the whole blood assay due to limited sample. The IFNγ response post-V1 and post-V2 was significantly higher than V0 healthy controls for both protein subunits (**Figure 2F**). The majority of the vaccinated samples tested using the whole blood assay were also positive for anti-spike antibodies (**Figure 2E)** and positive for T-cell responses to spike peptide pools in the ELISpot assay. Interestingly, one donor (P002) was positive for T-cell response using the whole blood and ELISpot assays, but negative for anti-spike IgG/A/M post-V1, and became positive for all three tests post-V2. Another subject initially in the control cohort (U002) later received the Moderna vaccine and was tested much earlier at 7 and 9 days post-V1. Positive T-cell responses to the spike protein were detected by both the whole blood and ELISpot assays, while remaining negative for a total antibody response until later **(Extended Data Figure 5**). These results demonstrated similar efficacy between the two tests (ELISpot and whole-blood assay) in identifying individuals with positive T-cell responses to SARS-CoV-2.

### Vaccine responses by demographics

We next interrogated the relationship of T-cell response in vaccinated subjects with time post vaccination, vaccine product, sex, and age (**Figure 3A**). To investigate if there was a difference in the kinetics and magnitude of the anti-spike T-cell response between individuals receiving either Moderna or Pfizer vaccines, we plotted the individual’s ELISpot SFU for spike pools A+B against time since vaccination (**Figure 3B**). In general, the median SFU increased following vaccination, including between post-V1 and post-V2. An example of one individual’s (M005) ELISpot shows an increasing response over the three time points (V0, post-V1, post-V2; **Figure 3C**). However, in three individuals who received the Moderna vaccine, the median SFU decreased between post-V1 and post-V2 assessment, though it still remained well above the 6.0 SFU threshold, possibly reflecting contraction of an effector T-cell response. One male subject vaccinated with Moderna (M019) was receiving the B-cell depleting drug rituximab at the time of vaccination and demonstrated an anti-spike T-cell response post-V1 (**Figure 3C**) while remaining negative for anti-spike IgG/A/M even post-V2, demonstrating the possibility of developing T-cell immunity independent of an antibody response. Overall, median SFUs post- V2 were increased in those receiving the Moderna vaccine compared to the Pfizer vaccine (spike pool A p=0.1069, spike pool B p=0.0805, A+B p=0.0596; **Figure 3D**). Across both vaccines, females had increased SFU for spike pool A+B compared to males (p=0.0411; **Figure 3E**). Age younger than 50 years was weakly associated with a higher median SFU response to spike pools A+B (p=0.2273; **Figure 3F** and **Extended Data Figure 6**).

### T-cell immunity to SARS-CoV-2 variants following vaccination

Using our standardized ELISpot assay, we also examined the impact of the emerging SARS-CoV- 2 variants on T-cell responses to the spike protein. By using individual peptide pools (rather than whole protein), our ELISpot assay provided sufficient dynamic range in T-cell responses to identify potential decrements in response to variant mutations. Variants such as B.1.1.7, B.1.351, and B.1.1.248/P1 were first detected in the United Kingdom, South Africa, and Brazil respectively. Not long after their discovery, these variants were detected globally and stirred concern due to their increased transmissibility and virulence^1–6^. There have also been reports of SARS-CoV-2 seropositive individuals becoming reinfected with a new variant^41–44^. These variants contain mutations in the spike protein (**Figure 4A**), including mutations in the ACE2 interacting surface of the RBD, that could lead to the loss of epitopes recognized by the neutralizing antibodies developed from SARS-CoV-2 infection and/or vaccination. In fact, multiple reports have demonstrated diminished neutralizing activity of vaccinated and convalescent patient sera in response to the variants compared to wild type SARS-CoV-2^10–12^.

Considering these variants may escape antibody neutralization, how they influence the T-cell immune response is of particular interest. One published report measured CD4 T-cell vaccine responses to a subset of the mutations in the B.1.1.7 and B.1.351 variants in one of two spike protein subunits, finding no decreased response to variant relative to wild type proteins^45^. However, as the authors acknowledge, wild type responses were very low due to the use of whole proteins rather than peptide pools^45^, which precludes an accurate fold-change assessment. To determine the impact of these variants on the magnitude of the anti-spike T-cell response in vaccinated individuals, we used our ELISpot assay to stimulate PBMC from vaccinated individuals post-V2 with spike peptide pools from the B.1.1.7, B.1.315, and B1.1.248 SARS-CoV-2 variants (**Figure 4A**). These pools were divided into pools A and B to mirror the wild type spike pools. Given the distribution of the mutations within the spike protein, the majority of the mutations, including those in the RBD, were present in spike pool A. There was a median decrease in SFU in response to spike pool A variants by 20.7% for B.1.1.7, 26.0% for B.1.351, and 41.4% for B.1.1.248 compared to the wild type. The response to spike pool B for B.1.1.7 and B.1.1.248 were not impacted (≤2.0% change); however, the median response to B.1.351 was decreased by 20.2% (**Figure 4B,C**). When combined, the response to variant spike pools A+B were decreased to 84.6% [95%CI 73.5–94.7] of the wild type response for variant B.1.1.7, 70.2% [62.3–86.7], for B.1.315, and 83.4% [58.3-97.0] for B.1.1.248 (**Figure 4D**). Additionally, there were several individuals who had a much greater reduction in their T-cell response to the variants despite having a good response to wild type peptides, suggesting that some individuals might have substantially impaired T-cell responses to the SARS-CoV-2 variants of interest. Median T-cell responses to the variants among donors receiving the Pfizer vaccine were lower than those receiving the Moderna vaccine, though not statistically significant (p=0.2446 for B.1.1.7, p=0.2455 for B.1.315, p=0.0740 for B.1.1.248; **Extended Data Figure 7**).

## DISCUSSION

Thus far in the course of the pandemic, seroconversion has been the focus of clinical tests of immunity, largely due to the ease of antibody detection. Unlike humoral immunity assays, measurements of T-cell immunity require functional viable cells, which makes these assays more challenging to deploy on a population-wide scale. Both T-cell immunity tests developed herein have excellent sensitivity and specificity and could be adopted by clinical laboratories to monitor the T-cell response to SARS-CoV-2 infection or vaccination using commercially available reagents. T-cell immunity may be especially important for convalescent individuals who do not seroconvert^32^ and immunocompromised patient populations who are less likely to develop an efficacious antibody response, as they may experience prolonged infection and/or longer duration of virus shedding^46^. In adults without effective humoral immunity as a result of underlying disease or therapeutic interventions, clinical measurements of T cell immunity may also guide physicians in determining when booster vaccination may be appropriate. Even in immunocompetent patients, increased T-cell response after SARS-CoV-2 infection has been associated with a more benign clinical course^47^. Our ELISpot assay could also help to distinguish between patients who were naturally infected with SARS-CoV-2 (positive response to the spike and nucleocapsid proteins) versus those who were vaccinated (response only to the spike protein) and can easily be modified to test for T-cell immunity to SARS-CoV-2 variants.

We used these assays to evaluate T-cell responses in convalescent and vaccinated individuals. A common gauge for vaccine potency is the development of antibodies to a similar level as those induced by natural infection. The initial SARS-CoV-2 vaccine trials demonstrated increases in IgG antibody levels and neutralizing activity following the full vaccination course that were similar to or higher than those observed in convalescent patients^36, 37^. Using this same parameter for T-cell responses, we found the number of SARS-CoV-2 spike protein-specific T-cells were increased in fully vaccinated individuals compared to convalescent patients who had experienced mild COVID-19. In our cohort, the T-cell responses increased significantly from baseline after the first dose of the vaccine (even as soon as 7 days after vaccination and before seroconversion), which was equivalent to responses in convalescent individuals. This is in contrast to reported antibody neutralization titers, which seem to require both doses to reach convalescent levels^36, 37^, unless the individual had prior SARS-CoV-2 exposure^48, 49^. The presence of a robust T-cell response comparable to convalescent patients after a single inoculation could suggest a similar level of SARS-CoV-2 protection and warrants further investigation as a possible measure to increase vaccine availability in resource-limited settings. Since duration of response will most likely be affected by the number of doses, long-term follow up of subjects receiving one or two doses will be valuable, including those receiving the Johnson & Johnson/Jansen vaccine who were not included in our study due to timing of FDA approval.

We observed a weak association between increased T-cell response in those vaccinated with Moderna compared to Pfizer. Though more subjects are needed to confirm the validity of this trend, such a difference could be a consequence of the higher dose of mRNA in the Moderna vaccine compared to Pfizer (100 μg and 30 μg respectively)^7, 8^. While no direct comparison of T-cell response between these two vaccines has previously been performed, a head-to-head comparison of the antibody responses found similar neutralizing activities^11^. We also identified a weak association between a decreased T-cell response and age, which has been similarly observed post-V2 with anti-spike IgG levels and antigen-specific memory B cells^48^. Finally, we observed an increased T-cell response in females compared to males, mirroring the response following natural infection where females with COVID-19 had more robust T-cell activation compared to males^50^.

The emergence of more transmissible and virulent SARS-CoV-2 variants^1–6^, coinciding with evidence of escape from neutralizing antibodies to the wild type virus^10–12^, raises concerns regarding the effectiveness of currently available vaccines. This doubt is highlighted by two recent cases of SARS-CoV-2 variant infection despite full vaccination in the presence of neutralizing antibodies to the original and variant viruses^51^. In the absence of effective humoral protection, T-cell immunity to these variants may be sufficient for viral clearance. We demonstrate reduced T-cell responses in vaccinated individuals to SARS-CoV-2 variants of concern B.1.1.7, B.1.351, and B.1.1.248. This reduction in T-cell response was predominantly to regions of the protein bearing the majority of the variant mutations, while responses to regions with fewer mutations were less affected. We observed a smaller fractional decrement in T-cell response to the variants than has been observed for antibody neutralization assays^10–12^. However, it will be important to determine how T-cell immunity impacts aggregate immune response to the variant SARS-CoV-2 viruses and if this reduction translates to adverse clinical outcomes.

## METHODS

### Study Participants

All donors were consented and participated voluntarily in this study. Protocols were approved by the Partners Institutional Review Board.

### Convalescent cohort

We tested 25 individuals (9 males, 16 females) who had recovered from COVID-19 between March and May 2020. The median age was 46 (range 25–69 years). SARS-CoV-2 infection was confirmed by PCR in 23/25 cases and by serology in the remaining 2/25 cases. Serology results for convalescent patients were extracted from medical records. The median time between the diagnostic result and the performance of the ELISpot assay was 162 days (83–237 days). The clinical symptoms reported for these donors were all classified as mild according to WHO definitions^1^. One donor was hospitalized briefly (C002).

### Healthy control cohorts

We tested 19 SARS-CoV-2 naive healthy controls (10 male, 9 female) by ELISpot. Ten donors were tested between the months of August and December 2020, with the remaining nine tested between January and April 2021. The median age 35.3 (23.0–63.7 years). Of the 19 donors, 13 had serology testing performed in parallel with the ELISpot for anti-spike IgG/A/M and all were negative. For the whole blood assay, we tested 7 healthy controls (4 male, 3 female, median age of 31.0, range 23.8–63.7). Healthy controls had no history of SARS-CoV-2 infection or close contact with a confirmed infected individual. All 7 donors had negative serology for anti-spike IgG/A/M.

### Vaccination cohorts

The ELISpot post-vaccination cohort included 14 donors (7 male, 7 female) after their first dose (2 Pfizer, 12 Moderna; median 22 days after the initial dose, range 16–30 days) with a median age of 35.6 (range 23.0–61.6 years). All 14 donors had serology testing performed for anti-spike IgG/A/M, of which only 2 did not seroconvert. An additional 29 donors (16 male, 13 female) were tested following their second dose (11 Pfizer, 18 Moderna; median 59 days after the initial dose, range 38–204 days) with a median age of 35.6 (23.0–81.0 years).

The post-V1 cohort for the whole blood assay included 12 donors (6 male, 6 female, 2 Pfizer, 10 Moderna, tested a median 22 days, range 18–30 days after vaccination), with a median age of 35.5 years (range 23.0–59.6 years). All 12 donors had serology for anti-spike IgG/A/M performed.

In addition, 16 donors (9 male, 7 female) were tested post-V2 (5 Pfizer, 11 Moderna, median 56 days, range 38–204), with a median age of 35.5 (range 23.0–73.3). One additional donor (U002) was only tested at day 7 and 9 post-V1 and is excluded from the vaccinated donor cohort and shown only in Extended Data Figure 5). Self-reported vaccine associated symptoms were collated and shown in **Extended Data Table 4**.

### Blood collection and processing

Blood was collected in Lithium Heparin tubes and red top serum tubes. Peripheral blood mononuclear cells (PBMC) were isolated from whole blood using ficoll density gradient centrifugation in a SepMate tube (Stem Cell Technologies) following 1:2 dilution with RPMI. Serum tubes were centrifuged, and serum was then aliquoted and stored at -80°C prior to performing the anti-spike antibody ELISA. ELISpot and whole blood assays were performed within 8 hours of venesection.

### Peptide pools and protein

The commercially available peptide pools consist of 15mers with an 11 amino acid overlap, spanning the entire spike and nucleocapsid sequence (JPT peptide solutions). Peptide pools based upon the initial sequence identified in Wuhan had a guaranteed purity of either >70% or >90%. Spike variant peptide pools are termed ‘crude’ with HPLC-MS assessment and chemical capping to prevent *de novo* epitope formation. The nucleocapsid pool (PM-WCPV-NCAP2) contains 102 peptides, whilst the spike protein pool (PM-WCPV-S-3) was divided into spike pool A (158 peptides) and spike pool B (157 peptides, of which the last is a 17mer). Spike pool A only contains peptides spanning the Spike S1 subunit from AA1-643, which includes the receptor binding domain, whilst spike pool B spans a small portion of the C terminal domain of S1 and the entire S2 (AA633-1273) (**Figure 1A**).

Peptide pools were reconstituted in 50uL DMSO and an equal volume sterile PBS (Gibco). Aliquots were stored at -20°C. Lot to lot comparisons were performed for different purity pools (>70% versus >90%) using SARS-Cov-2 vaccinated donors.

### IFN**γ** ELISpot assay

PBMC were resuspended in serum free T-cell assay media (ImmunoSpot) to give a concentration of 2.5x10^6^/mL and 100 μL was added to each well of the Human IFNγ single color ELISpot plate (ImmunoSpot). Cells were incubated with 1:1 DMSO:PBS (negative control); the spike A, spike B, or nucleocapsid peptide pools; or phytohemagglutinin (PHA, positive control). The controls and antigens were added in 50 μL to each well, where possible in duplicate wells. PHA (Sigma-Aldrich) was added as a positive control at a final concentration of 5 μg/mL. Peptide pools contained each peptide at a final concentration of 2 μg/mL. Negative control wells contained the equivalent volume of DMSO and PBS as the peptide pools. Peptide pools were thawed immediately prior to use. The plate was incubated at 37°C for 16-20 hours. Plates were developed according to manufacturer’s instructions and air dried before counting on an ImmunoSpot CoreS6 ELISpot counter (ImmunoSpot). QC of automated count data was performed by a lab member not involved in assay set up to minimize operator bias. The background SFU count was subtracted from the antigen wells spot count and the average SFU result for spike pool A, spike pool B, and nucleocapsid pool are reported. In some instances, the responses to spike pool A and B are added together to better reflect responses to the entire spike protein. In cases where there were no spot responses to antigen, the PHA well was always strongly positive. The median background in the negative control was 0 SFU/2.5x10^5^ cells (range 0-6). The positive threshold of 6 SFU per 2.5x10^5^ PBMC was determined via receiver operator characteristic (ROC) curve analysis.

### Selection of ELISpot SFU threshold for positivity

Receiver operator characteristic (ROC) curve analysis was performed in Prism V8.0 (GraphPad Software Inc) using the unvaccinated donors with no known history of SARS-CoV-2 infection or contact with known infected individuals (healthy controls), alongside the 25 convalescent patients. This was performed individually for spike peptide pool A, spike peptide pool B and the nucleocapsid peptide pool (**Extended Data Figure 2**). The impact of altering the mean SFU assay threshold on clinical sensitivity and specificity was evaluated when utilizing the responses to either spike pool A or B. The nucleocapsid result was not included with the spike responses as a response to the nucleocapsid would not be induced to the current spike vaccines.

### Whole blood assay and IFN**γ** ELISA

Overnight incubations with negative control, positive control, S1 and S2 proteins were performed from donors at three time points (V0, post-V1, and post-V2). Peripheral blood was mixed thoroughly before aliquoting 1 mL into five sterile tubes with loose fitting lids. For the negative condition, 20 μL of sterile water is added to the tube. For positive controls, 18 μL of PHA (1 μg/μL stock) and 2 μL of Cell activation cocktail (Biolegend) are added to the tube. The whole protein spike S1 subunit, spike S2 subunit of SARS-CoV-2 are thawed just prior to use and added in 20 μL to individual tubes to give a final concentration of 5 mg/mL. Samples are carefully and thoroughly mixed before incubating at 37°C for 16–24 hours. Samples were then centrifuged and plasma isolated and stored at -20°C prior to testing in the QuantiFERON IFNγ ELISA (Qiagen). Acceptance criteria for the assay specified by manufacturers were always met.

### Selection of IFN**γ** ELISA IU/ml threshold for positivity

ROC curve analysis was performed in Prism V8.0 (GraphPad Software Inc) for IFNγ release against S1 and the S2 proteins using 7 unvaccinated healthy donors with no known history of SARS-CoV-2 infection or close contact with known infected individuals, alongside 16 fully vaccinated donors (11 Moderna and 5 Pfizer). A threshold of ≥0.3 IU/mL IFNγ was selected. The combined use of IFNγ results from both S1 and S2 proteins was evaluated to determine the sensitivity and specificity of the combination assay using a confusion matrix. The upper limit of detection is 10 IU/mL and samples with ≥10 IU/mL were not further diluted.

### ELISA for detection of antibody response to SARS-CoV-2 spike protein

A qualitative ELISA for Human Anti-IgG/A/M SARS-CoV-2 ELISA (The Binding Site) was performed using donor serum per manufacturer’s instructions. Serum was diluted 1:40 prior to the assay. Plates were read at 450nm using a BioTek Synergy microplate reader. A cut-off calibrator and a high and a low control were run in each assay. Results were calculated using the equation: Calibrator coefficient x (Sample OD / Cut-off calibrator OD). A result of <1.0 is interpreted as negative for anti-spike antibodies, and a result of ≥1.0 is interpreted as positive for anti-spike antibodies. The clinical specificity of this assay is reported to be 98.4% [95% CI, 96.6–99.3%], and sensitivity 94.7% [95% CI, 90.9–97.2%].^2^

### Variant modeling

Spike protein was modeled using Pymol version 2.4.1 and the protein data bank trimeric spike protein structure 7DX0 in which one of the spike monomers’ receptor binding domains adopts the “up” conformation^3^. In some cases, specific spike variant residues were not modeled in the original structure and adjacent residues were highlighted as representative. In all protein sequence representations, the abbreviations are as follows. Spike: Ss, single sequence; NTD, N terminal-domain; RBD, receptor binding domain; SD1-2, subdomain 1 subdomain 2; FP, fusion peptide; HR1, heptad repeat 1; CH, central helix; CD, connector domain; HR2, heptad repeat 2; TM, transmembrane domain; IC, intracellular domain. Nucleocapsid protein: NTD, N terminal domain; RBD, RNA binding domain; CTD, C terminal domain; DD, dimerization domain.

### Statistical analysis

Statistical analysis was performed using Prism V8.0 (GraphPad Software Inc). Normality of data was evaluated using the Shapiro-Wilk test. Log transformation was performed where appropriate for non-normally distributed data. Where distribution remained non-normal, non-parametric tests were performed. Parametric tests were performed where data was normally distributed. Evaluation of differences between matched pairs was performed using the Wilcoxon matched pairs signed-ranks test and the Bonferroni correction for multiple comparisons. The differences between unmatched groups were compared using the Kruskal-Wallis test, the Mann Whitney U test or a paired t test. For analysis of ELISpot responses to variant pools, only responses for which there were ≥6 SFU were included in the statistical analysis.

## ACKNOWLEDGEMENTS

This work was conducted with biostatistical support from Harvard Catalyst and R01CA238268, R01CA252940, R01CA249062 (MVM). Zachary J Manickas Hill for evaluating convalescent patient medical records. Thank you to Tamina Kienka, Andrea Schmidts, Jane O, and Madeline Polak for phlebotomy, and Emily Silva and Max Jan for transporting samples. The MGH/MassCPR COVID biorepository was supported by a gift from Ms. Enid Schwartz, by the Mark and Lisa Schwartz Foundation, the Massachusetts Consortium for Pathogen Readiness and the Ragon Institute of MGH, MIT and Harvard. We would also like to thank all the participants for volunteering in the study.

## AUTHOR CONTRIBUTIONS

K.M.E.G. and M.V.M. conceived and designed the study. K.M.E.G., K.K., J.Y.Y., G.B., and performed the experiments. MGH COVID-19 Collection and Processing Team provided convalescent patient samples. K.G. counted the ELISpots. K.M.E.G., M.B.L., and R.C.L. analyzed data. K.M.E.G., M.B.L., R.C.L., T.R.B., and M.V.M. wrote the paper with help from all authors.

## COLLABORATORS

### MGH COVID-19 Collection & Processing Team

**Collection Team:** Kendall Lavin-Parsons^1^, Blair Parry^1^, Brendan Lilley^1^, Carl Lodenstein^1^, Brenna McKaig^1^, Nicole Charland^1^, Hargun Khanna^1^, Justin Margolin^1^

**Processing Team:** Anna Gonye^2^, Irena Gushterova^2^, Tom Lasalle^2^, Nihaarika Sharma^2^, Brian C. Russo^3^, Maricarmen Rojas-Lopez^3^, Moshe Sade-Feldman^4^, Kasidet Manakongtreecheep^4^, Jessica Tantivit^4^, Molly Fisher Thomas^4^

**Massachusetts Consortium on Pathogen Readiness:** Betelihem A. Abayneh^5^, Patrick Allen^5^, Diane Antille^5^, Katrina Armstrong^5^, Siobhan Boyce^5^, Joan Braley^5^, Karen Branch^5^, Katherine Broderick^5^, Julia Carney^5^, Andrew Chan^5^, Susan Davidson^5^, Michael Dougan^5^, David Drew^5^, Ashley Elliman^5^, Keith Flaherty^5^, Jeanne Flannery^5^, Pamela Forde^5^, Elise Gettings^5^, Amanda Griffin^5^, Sheila Grimmel^5^, Kathleen Grinke^5^, Kathryn Hall^5^, Meg Healy^5^, Deborah Henault^5^, Grace Holland^5^, Chantal Kayitesi^5^, Vlasta LaValle^5^, Yuting Lu^5^, Sarah Luthern^5^, Jordan Marchewka (Schneider)^5^, Brittani Martino^5^, Roseann McNamara^5^, Christian Nambu^5^, Susan Nelson^5^, Marjorie Noone^5^, Christine Ommerborn^5^, Lois Chris Pacheco^5^, Nicole Phan^5^, Falisha A. Porto^5^, Edward Ryan^5^, Kathleen Selleck^5^, Sue Slaughenhaupt^5^, Kimberly Smith Sheppard^5^, Elizabeth Suschana^5^, Vivine Wilson^5^, Galit Alter^6^, Alejandro Balazs^6^, Julia Bals^6^, Max Barbash^6^, Yannic Bartsch^6^, Julie Boucau^6^, Josh Chevalier^6^, Fatema Chowdhury^6^, Kevin Einkauf^6^, Jon Fallon^6^, Liz Fedirko^6^, Kelsey Finn^6^, Pilar Garcia-Broncano^6^, Ciputra Hartana^6^, Chenyang Jiang^6^, Paulina Kaplonek^6^, Marshall Karpell^6^, Evan C. Lam^6^, Kristina Lefteri^6^, Xiaodong Lian^6^, Mathias Lichterfeld^6^, Daniel Lingwood^6^, Hang Liu^6^, Jinqing Liu^6^, Natasha Ly^6^, Ashlin Michell^6^, Ilan Millstrom^6^, Noah Miranda^6^, Claire O’Callaghan^6^, Matthew Osborn^6^, Shiv Pillai^6^, Yelizaveta Rassadkina^6^, Alexandra Reissis^6^, Francis Ruzicka^6^, Kyra Seiger^6^, Libera Sessa^6^, Christianne Sharr^6^, Sally Shin^6^, Nishant Singh^6^, Weiwei Sun^6^, Xiaoming Sun^6^, Hannah Ticheli^6^, Alicja Trocha-Piechocka^6^, Daniel Worrall^6^, Alex Zhu^6^, George Daley^7^, David Golan^7^, Howard Heller^7^, Arlene Sharpe^7^, Nikolaus Jilg^8^, Alex Rosenthal^8^, Colline Wong^8^

^1^Department of Emergency Medicine, Massachusetts General Hospital, Boston, MA, USA. ^2^Massachusetts General Hospital Cancer Center, Boston, MA, USA. ^3^Division of Infectious Diseases, Department of Medicine, Massachusetts General Hospital, Boston, MA, USA. ^4^Massachusetts General Hospital Center for Immunology and Inflammatory Diseases, Boston, MA, USA. ^5^Massachusetts General Hospital, Boston, MA, USA. ^6^Ragon Institute of MGH, MIT and Harvard, Cambridge, MA, USA. ^7^Harvard Medical School, Boston, MA, USA. ^8^Brigham and Women’s Hospital, Boston, MA, USA.

## ADDITIONAL INFORMATION

Correspondence and requests for materials should be addressed to: Marcela Maus

## EXTENDED DATA

**Extended Data Table 1.** Healthy controls used to establish ELISpot and whole blood assay. Donors had no history of infection with SARS-CoV-2 or close contact with confirmed SARS-CoV-2 infected individuals. 19 donors were tested in the ELISpot assay and 7 donors were tested in the whole blood assay. 15 donors had serum tested for the presence of anti-spike IgG/A/M by ELISA. Where an assay was not performed the box is grey filled. Results are reported as positive (P) or negative (N) according to the final assay thresholds.

| Donor ID | Age (Years) | Sex (M-male, F-female) | Anti-spike IgG/A/M | IFNγ ELISpot response to (spike pools A/B) | Whole blood assay result (S1, S2 protein) |
| --- | --- | --- | --- | --- | --- |
| M002 | 39.7 | F | N | N |  |
| M003 | 35.6 | F | N | N |  |
| M004 | 35.3 | M |  | P |  |
| M005 | 24.9 | M | N | N |  |
| M007 | 36.2 | M |  | P |  |
| M008 | 23.0 | F |  | N |  |
| M009 | 27.6 | F |  | N |  |
| M011 | 45.0 | F |  | N |  |
| M012 | 50.8 | M | N | N |  |
| M013 | 25.8 | M | N | N |  |
| P001 | 33.9 | M | N | N |  |
| P000 | 63.7 | F | N | N | N |
| U001 | 33.5 | F |  | N |  |
| U002 | 27.3 | M | N | N | N |
| U003 | 28.0 | M | N | N | N |
| U004 | 35.6 | F | N | N | N |
| U005 | 36.1 | M | N | N | N |
| U006 | 59.5 | F | N | N |  |
| U007 | 33.7 | M | N | N |  |
| U009 | 31.0 | M | N |  | N |
| U010 | 23.8 | F | N |  | N |

**Extended Data Table 2.** Convalescent patients used to establish ELISpot assay. Donors with previously confirmed COVID-19 from which they recovered. Diagnosis was confirmed either by PCR or antibody testing. Data was provided from medical records. Where an assay was not performed the box is grey filled. Results are reported as positive (P) or negative (N) according to the final assay thresholds.

| Patient ID | Patient Age (Years) | Sex (M-male, F-female) | Hospitalised for COVID (Y-Yes, N-No) | PCR result | Antibody Result | Days since COVID-19 diagnostic test | ELISpot response to spike pools | ELISpot response to NC pool |
| --- | --- | --- | --- | --- | --- | --- | --- | --- |
| C001 | 57 | F | N | P |  | 126 | P | P |
| C002 | 62 | F | Y | P |  | 164 | P | N |
| C003 | 69 | M | N | P |  | 134 | P | P |
| C004 | 60 | F | N | N | P (IgG, IgM) | 95 | P | N |
| C005 | 66 | F | N | P |  | 226 | P | P |
| C006 | 28 | M | N | P |  | 125 | P | P |
| C007 | 37 | F | N | P |  | 130 | P | P |
| C008 | 46 | F | N | N | P (IgM) | 83 | N | N |
| C009 | 26 | M | N | P |  | 209 | P | P |
| C010 | 29 | F | N | P |  | 210 | P | P |
| C011 | 46 | F | N | P | N | 235 | N | P |
| C012 | 69 | F | N | P |  | 204 | P | P |
| C013 | 69 | M | N | P |  | 204 | P | P |
| C014 | 30 | F | N | P |  | 103 | P | P |
| C015 | 25 | F | N | P |  | 111 | P | P |
| C016 | 28 | M | N | P |  | 237 | P | P |
| C017 | 53 | M | N | P |  | 85 | P | N |
| C018 | 47 | F | N | P |  | 134 | P | P |
| C019 | 32 | M | N | P |  | 223 | P | P |
| C020 | 31 | F | N | P |  | 230 | P | P |
| C021 | 62 | F | N | P |  | 204 | P | P |
| C022 | 32 | F | N | P |  | 231 | P | P |
| C023 | 34 | F | N | P |  | 109 | P | P |
| C024 | 64 | M | N | P |  | 162 | P | P |
| C025 | 49 | M | N | P |  | 131 | P | P |

**Extended Data Table 3.**
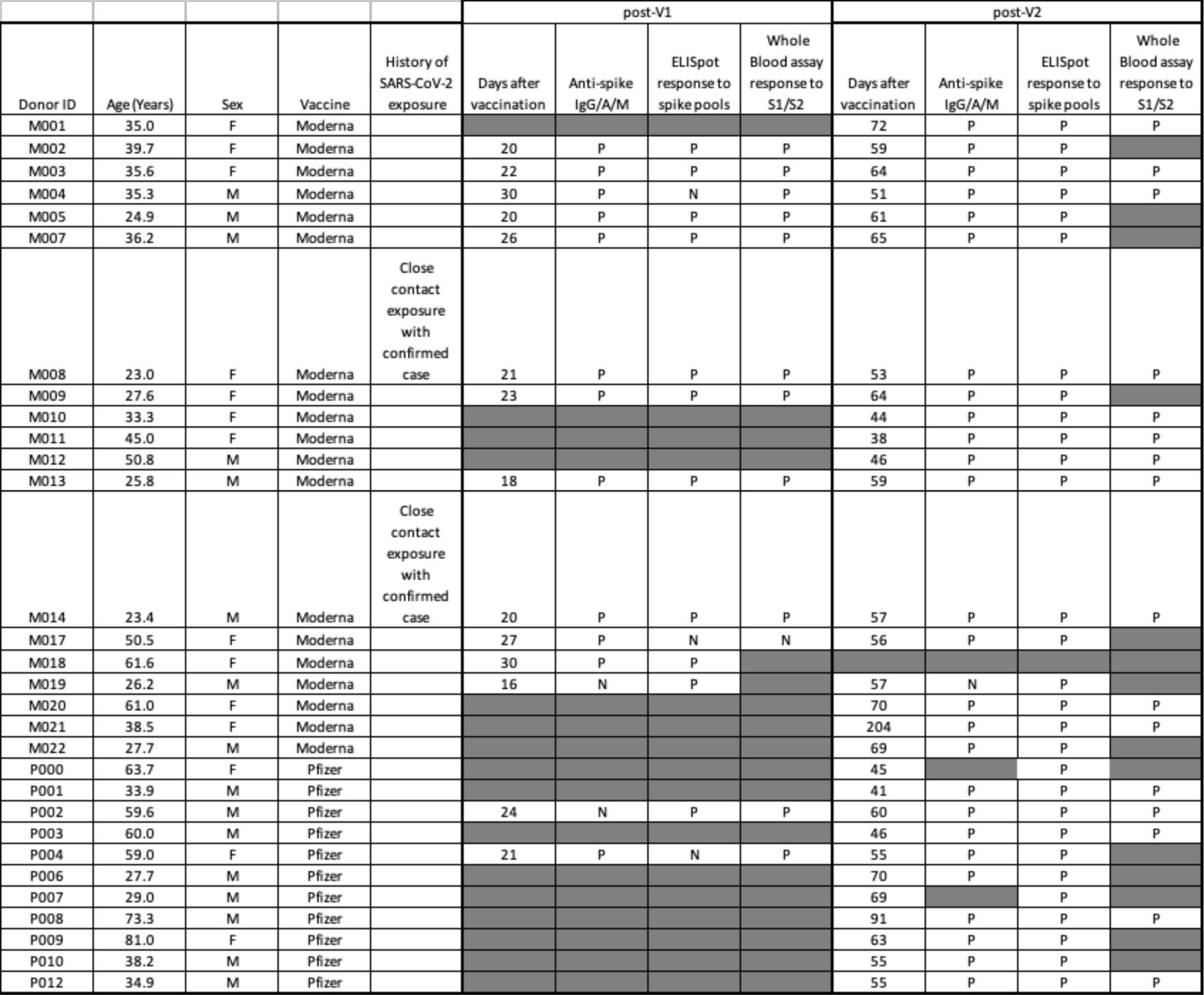
Vaccinated donors tested at V1 and V2 in ELISpot and whole blood assays. Donors tested at V1 and/or V2 for T cell responses to spike using the ELISpot assay and/or the whole blood assay. Where an assay was not performed the box is grey filled. Results are reported as positive (P) or negative (N) according to the final assay thresholds. No donors had confirmed SARS-CoV-2 infection. Instances of close contact exposure with a confirmed infected individual were documented.

**Extended Data Table 4.** Vaccinated donor post-vaccine symptoms. All vaccinated donors provided self reported symptoms in the days following vaccination.

| Donor | Age (years) | Sex | Vaccine | No Symptoms | Pain and swelling in arm | Fever or feeling feverish | Fatigue | Myalgia (muscle aches) | Other |
| --- | --- | --- | --- | --- | --- | --- | --- | --- | --- |
| M001 | 35.0 | F | Moderna |  | x |  | x |  |  |
| M002 | 39.7 | F | Moderna |  | x | x | x |  |  |
| M003 | 35.6 | F | Moderna |  |  |  | x | x |  |
| M004 | 35.3 | M | Moderna |  | x | x | x |  |  |
| M005 | 24.9 | M | Moderna |  |  | x | x |  |  |
| M007 | 36.2 | M | Moderna |  | x | x | x |  |  |
| M008 | 23.0 | F | Moderna |  | x | x | x |  |  |
| M009 | 27.6 | F | Moderna |  | x | x | x | x |  |
| M010 | 33.3 | F | Moderna |  | x |  | x | x | Headache |
| M011 | 45.0 | F | Moderna |  | x |  | x |  |  |
| M012 | 50.8 | M | Moderna |  |  | x |  | x |  |
| M013 | 25.8 | M | Moderna |  | x |  |  |  |  |
| M014 | 23.4 | M | Moderna |  | x | x | x | x | Headache |
| M017 | 50.5 | F | Moderna |  | x | x | x | x |  |
| M018 | 61.6 | F | Moderna |  | x |  |  |  |  |
| M019 | 26.2 | M | Moderna |  |  |  | x |  |  |
| M020 | 61.0 | F | Moderna |  | x |  | x | x |  |
| M021 | 38.5 | F | Moderna |  | x | x | x | x | pre-syncope |
| M022 | 27.7 | M | Moderna |  | x |  |  |  |  |
| P000 | 63.7 | F | Pfizer | x |  |  |  |  |  |
| P001 | 33.9 | M | Pfizer | x |  |  |  |  |  |
| P002 | 59.6 | M | Pfizer |  | x |  |  |  |  |
| P003 | 60.0 | M | Pfizer |  | x |  |  | x |  |
| P004 | 59.0 | F | Pfizer | x |  |  |  |  |  |
| P006 | 27.7 | M | Pfizer |  |  |  |  |  | Headache |
| P007 | 29.0 | M | Pfizer |  |  |  | x |  |  |
| P008 | 73.3 | M | Pfizer |  | x |  | x |  |  |
| P009 | 81.0 | F | Pfizer |  |  |  | x |  |  |
| P010 | 38.2 | M | Pfizer | x |  |  |  |  |  |
| P012 | 34.9 | M | Pfizer |  | x |  |  |  |  |

**Extended Data Figure 1.**
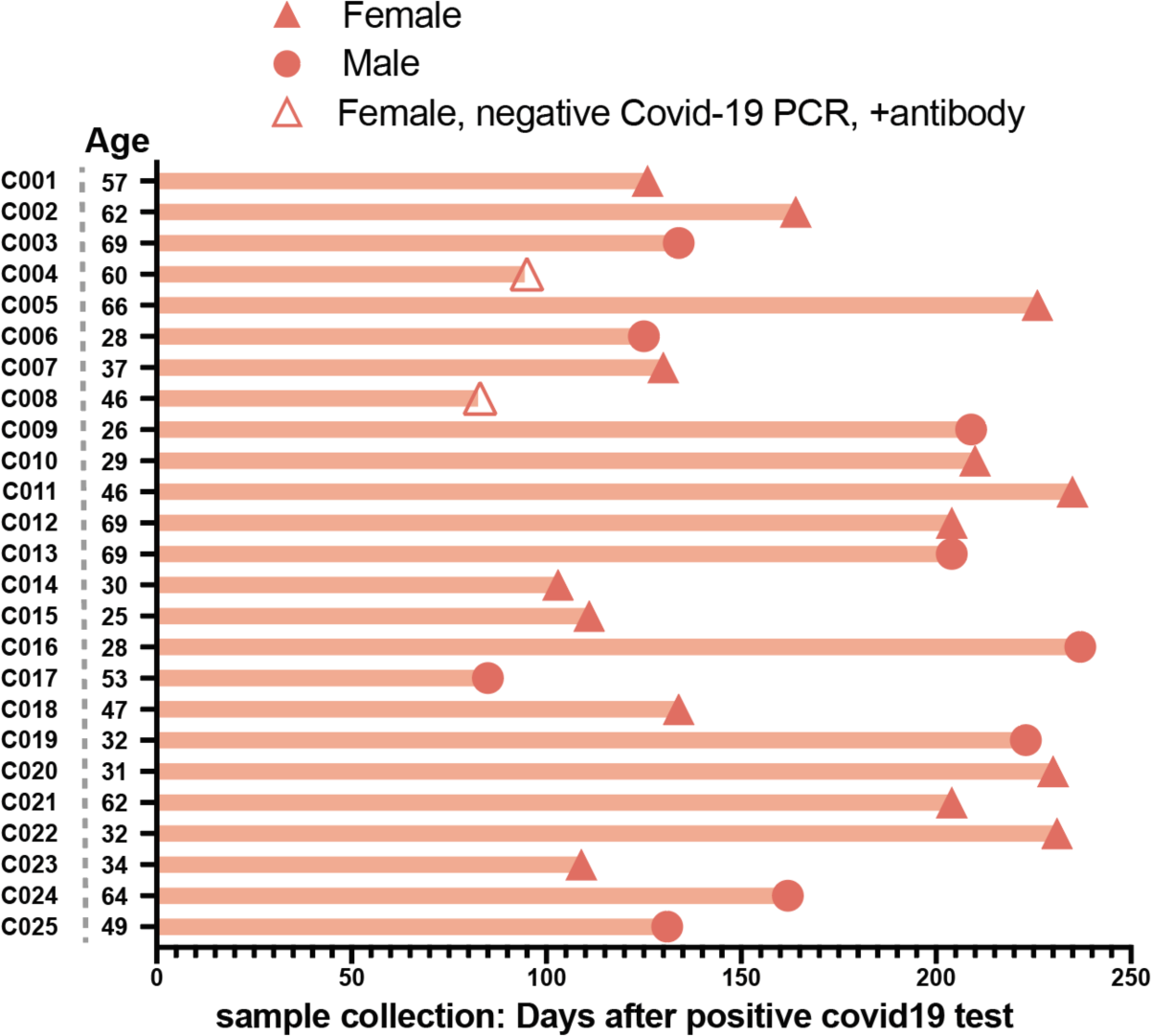
Convalescent patient cohort characteristics. n=25. 9 males and 16 females who had recovered from COVID-19 between March and May 2020. Diagnosis was based upon PCR testing and/or antibody testing.

**Extended Data Figure 2.**
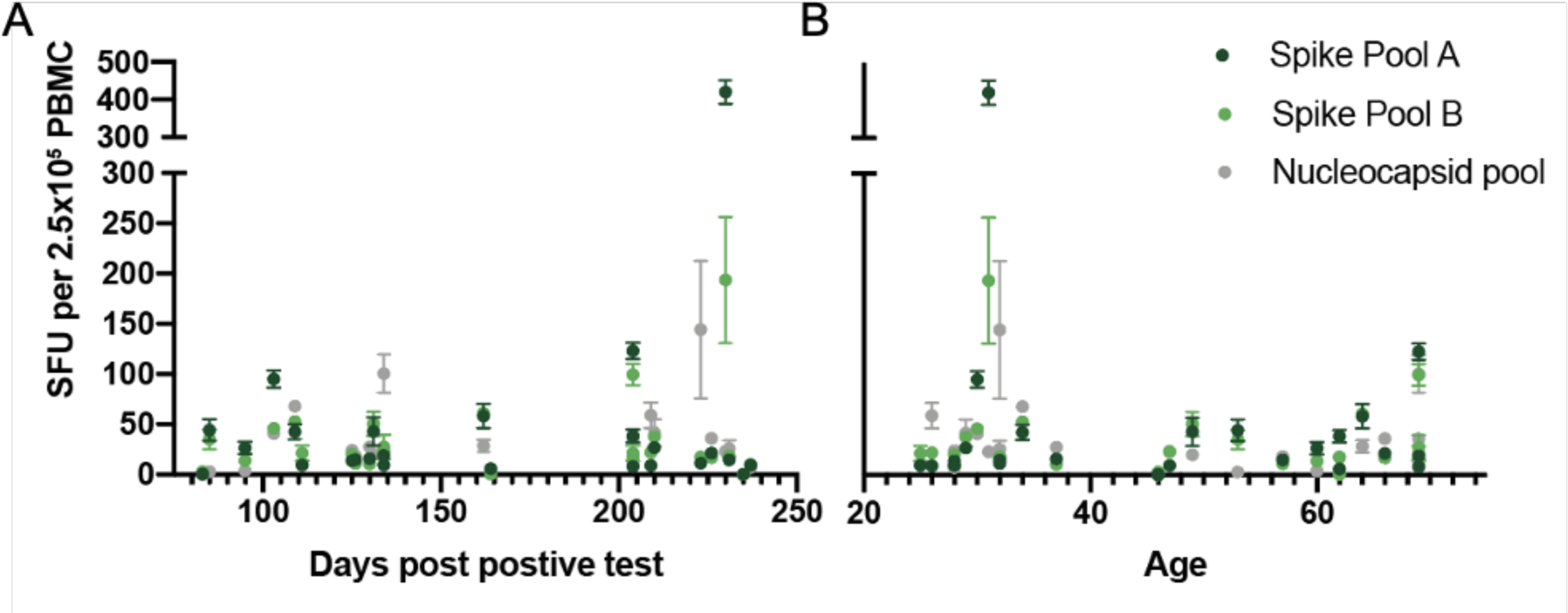
Convalescent patient cohort responses by sample collection timing and age. **A.** Simple linear regression between days post positive test and spike pool A resulted in an R^2^ of 0.0382, spike pool B R^2^=0.0238, and the nucleocapsid pool R^2^=0.0258. **B.** Linear regression between age and spike pool A resulted in an R^2^ of 0.0130, spike pool B R^2^=0.0508, and the nucleocapsid pool R^2^=0.0185.

**Extended Data Figure 3.**
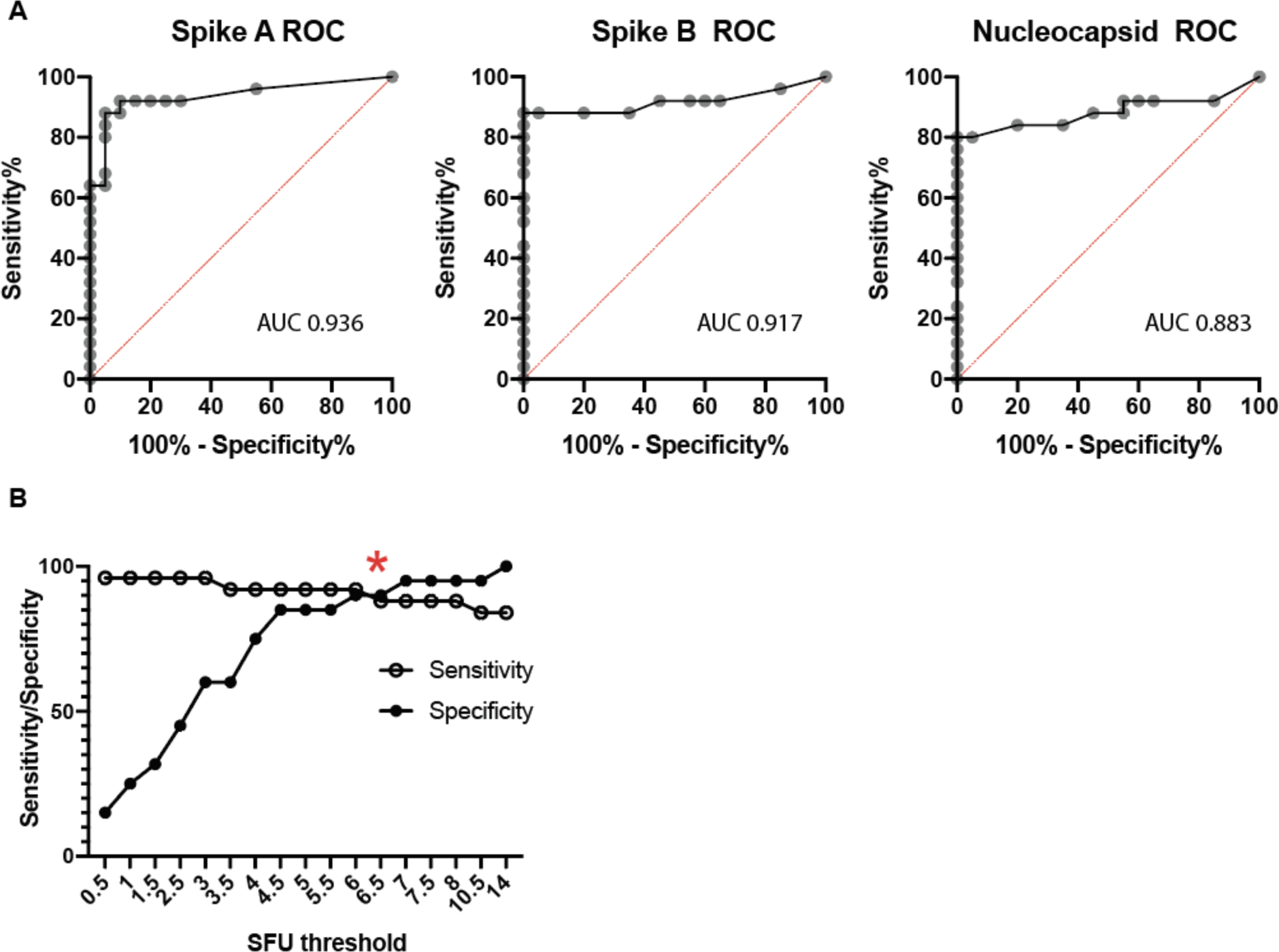
Test characteristics for the ELISpot SARS CoV-2 assay. **A**. Receiver operator characteristic (ROC) curves for individual results for spike pool A, B, and the nucleocapsid peptide pools utilizing data from the convalescent and healthy donor subjects in Figure 1. Area under the curve (AUC) is shown for each ROC. **B**. Evaluation of SFU positive threshold for responses to either spike A or to spike B pools as a combined output versus sensitivity and specificity. Red asterisk represents the selected threshold to optimize the tradeoff between sensitivity and specificity. Convalescent patients had a median SFU of 16.0, 95% CI [9.5–38.5] for spike pool A, 21.5 [17.0–35.0] for spike pool B, and 23.5 [12.0–36.0] for the nucleocapsid pool, compared to 0.5 [0.5–1.5], 2.5 [0.5–3.5], and 0.0 [0.0–1.0] in the healthy controls.

**Extended Data Figure 4.**
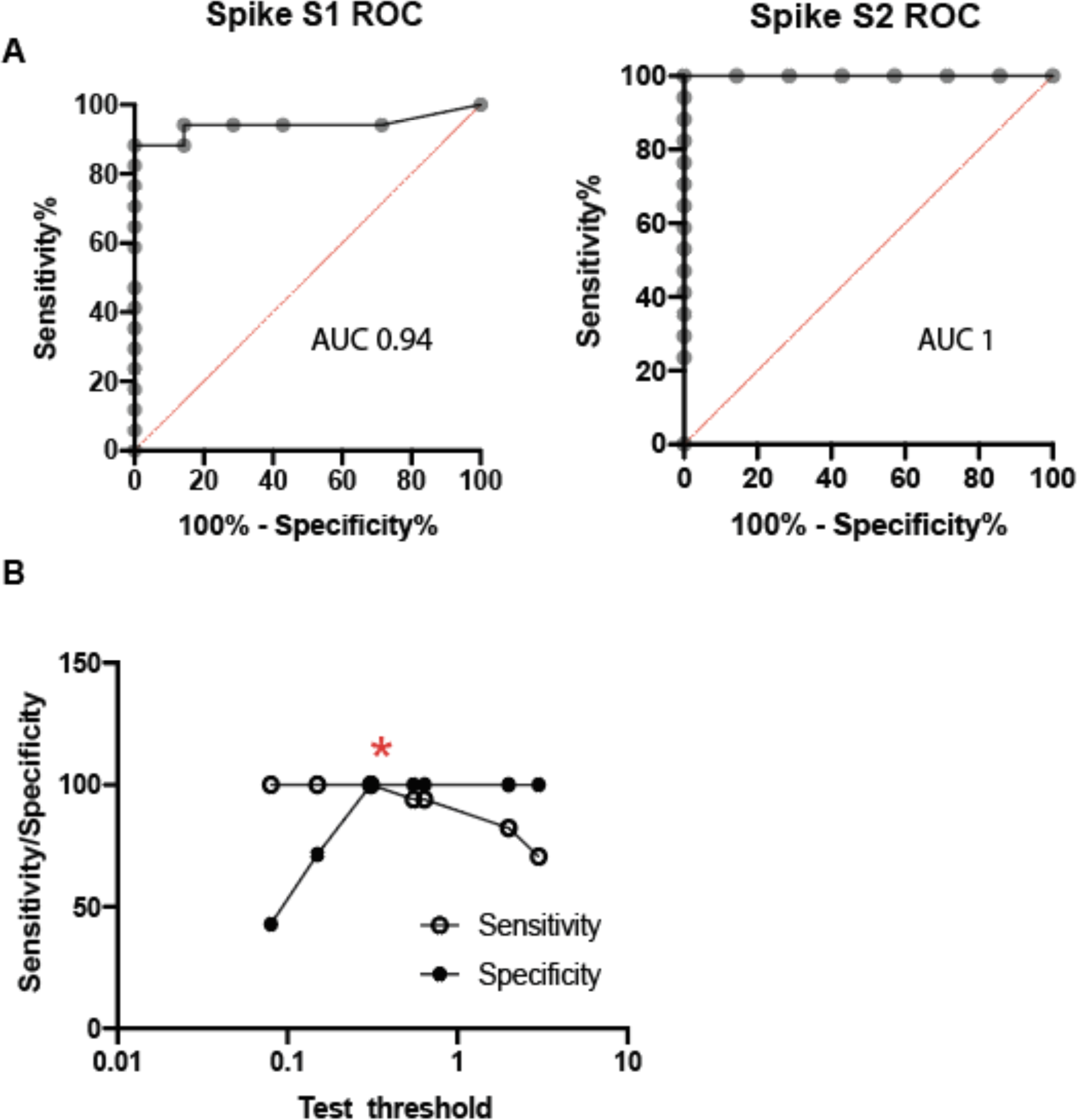
Test characteristics for the whole blood quantiferon assay. **A.** Receiver operator characteristic (ROC) curves for individual results for spike subunit 1 and subunit 2. Area under the curve (AUC) is shown for each ROC. **B.** Evaluation of IFNγ positive threshold for responses to either spike subunit 1 or 2 as a combined output versus sensitivity and specificity. Red asterisk represents the selected threshold to optimize the tradeoff between sensitivity and specificity.

**Extended Data Figure 5.**
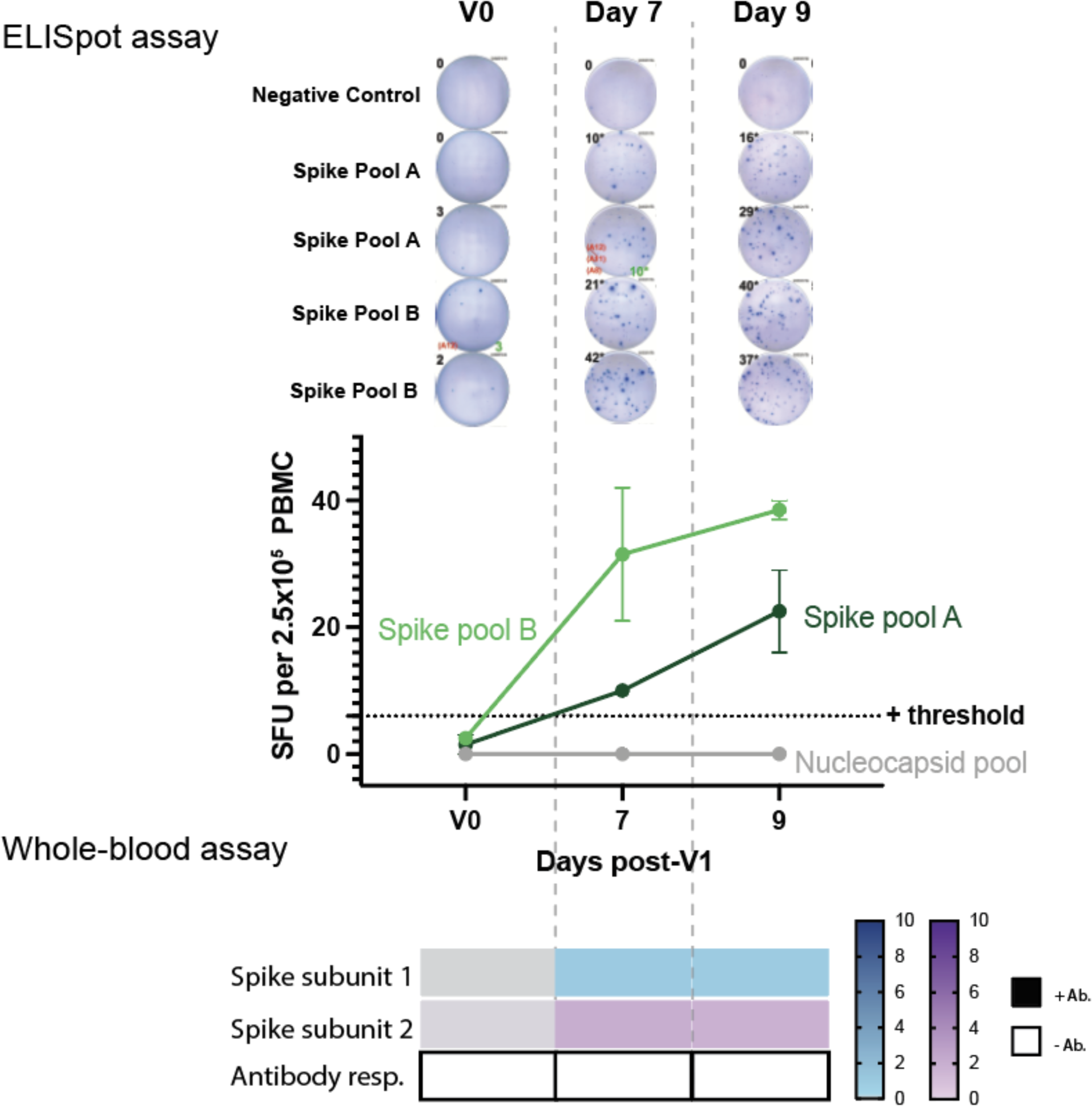
Early detection of SARS-CoV-2 T cell response post-V1. Healthy control subject U002 underwent evaluation via ELISpot assay and whole-blood assay before vaccination (V0) as well as 7 and 9 days after the first dose of the Moderna vaccine. Total antibody response (IgA/IgG/IgM) was negative at these time points and later seroconverted.

**Extended Data Figure 6.**
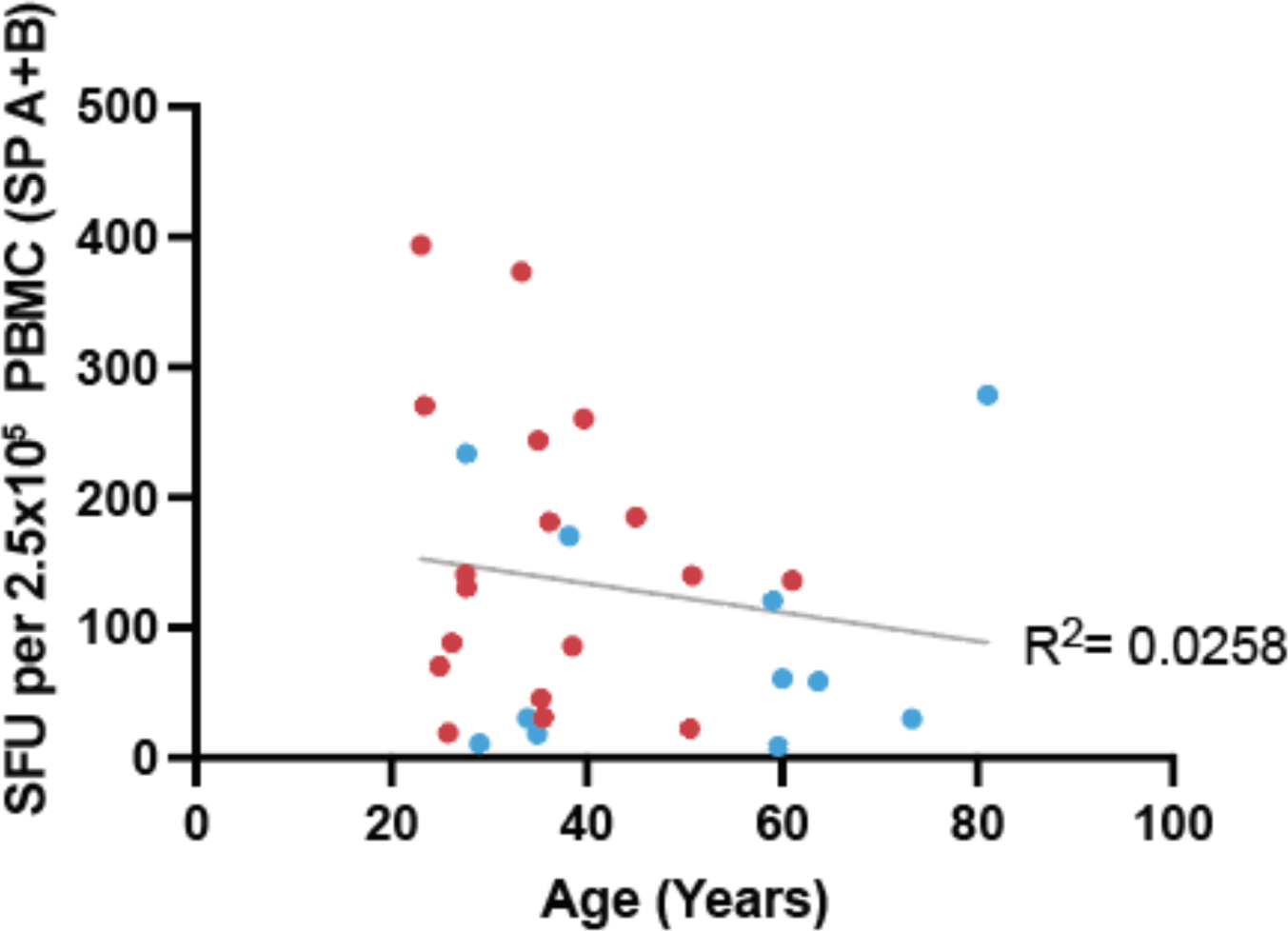
Spike pool A+B response as a function of age. Sum of the mean ELISpot results for spike pool A and B are shown as a function of age. R^2^ represents the goodness of fit for a simple linear regression. Moderna (red) and Pfizer (blue).

**Extended Data Figure 7.**
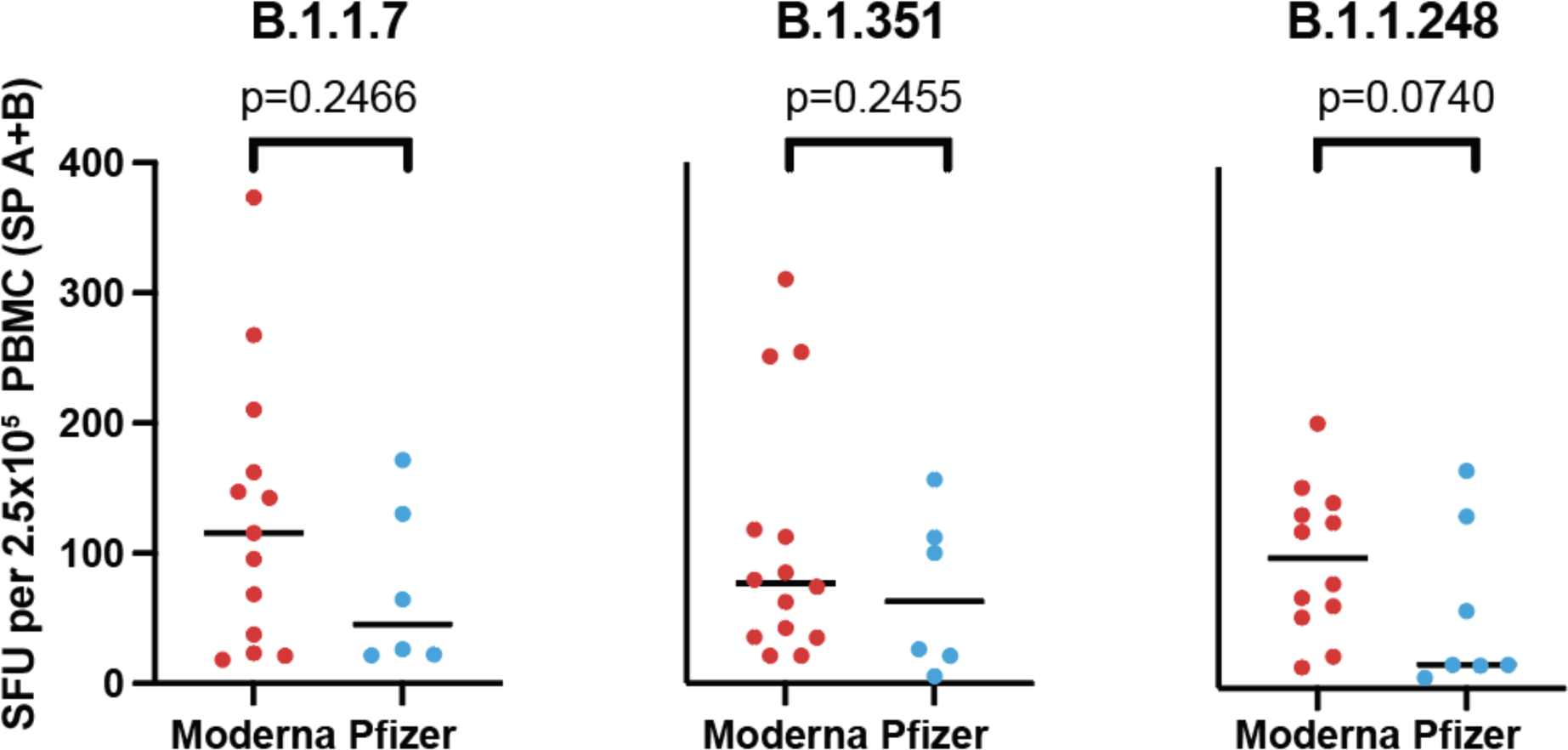
Variant response separated by vaccine. Sum of the mean ELISpot results for spike pool A and B are shown for each tested variant stratified by type of vaccine the subject received. Comparisons were performed with individual unpaired t-tests.

